# Hypercluster: a python package and SnakeMake pipeline for flexible, parallelized unsupervised clustering optimization

**DOI:** 10.1101/2020.01.13.905323

**Authors:** Lili Blumenberg, Kelly V. Ruggles

**Affiliations:** Institute of Systems Genetics, New York University School of Medicine, New York, NY 10016, USA; Department of Medicine, New York University School of Medicine, New York, NY 10016, USA

## Abstract

Unsupervised clustering is a common and exceptionally useful tool for large biological datasets. However, clustering requires upfront algorithm and hyperparameter selection, which can introduce bias into the final clustering labels. It is therefore advisable to obtain a range of clustering results from multiple models and hyperparameters, which can be cumbersome and slow. To streamline this process, we present hypercluster, a python package and SnakeMake pipeline for flexible and parallelized clustering evaluation and selection. Hypercluster is available on bioconda; installation, documentation and example workflows can be found at: https://github.com/ruggleslab/hypercluster.

**Author summary:** Unsupervised clustering is a technique for grouping similar samples within a dataset. It is extremely common when analyzing big data from patient samples, or high throughput techniques like single cell RNA-seq. When researchers use unsupervised clustering, they have to select parameters that affect the final result—for instance, how many groups they expect to find or what the smallest group is allowed to be. Some methods require setting even less intuitive parameters. For most applications, it is extremely challenging to guess what the values of these parameters should be; therefore to prevent introducing bias into the final results, researchers should test many different parameters and methods to find the best groups. This process is cumbersome, slow and challenging to perform in a reproducible way. We developed hypercluster, a tool that automates this process, make it much faster, and presenting the results in a reproducible and helpful manner.

## Introduction

Unsupervised clustering is commonly used for the interpretation of ‘omics datasets. It provides an objective and intuitive measure of similarity and difference between samples. Clustering can be used to determine biologically relevant subgroups of samples, find co-regulated molecular features, or provide objective support for the phenotypic similarity of biological perturbations. Moreover, clustering is a key step in the analysis of many emerging sequencing-based technologies. For example, a fundamental challenge in the analysis of single-cell measurement data, in particular single cell RNA-seq (scRNA-seq), is determining robust clusters of phenotypically similar cells (1–3). Clustering is also increasingly being used alongside traditional diagnostic techniques to establish new classifications of patient samples into disease-relevant subgroups (4–7) and for patient subgroup classification and risk stratification (6,8–12). The near-future of personalized medicine relies on researchers identifying robust unsupervised clustering-based disease subtypes. Therefore, it is essential that high-quality clustering results are easily and robustly obtainable, without user-selected hyperparameters introducing bias and impeding rapid analysis.

Currently, researchers robustly employing unsupervised clustering must choose specific algorithms and hyperparameters that are appropriate to their experiment type and data. Although some efforts have been made to advise researchers on optimal selection of both (13), biological datasets vary between batches, days, labs and researchers, underlining the importance of context- and experiment-dependent analysis tuning. Software packages for automatic hyperparameter tuning and model selection for regression and classification machine learning techniques exist, notably auto-sklearn from AutoML (14), but there are not yet packages for automated unsupervised clustering optimization.

Typically, the effect of hyperparameter choice on the quality of clustering results cannot be described with a convex function, meaning that when searching the landscape of hyperparameter choices there are often local maxima that may appear to be the optimal results if broad choices of hyperparameters are not considered. Therefore it is unlikely that a sequential approach using for instance, gradient descent from a single initialized set of hyperparameters, would be able to select the optimal parameters for the majority of clustering challenges (15). Exhaustive (i.e. grid) search is the most likely to obtain optimal results from unsupervised clustering. However, grid search can be slow and cumbersome to perform for the multiple hyperparameters and clustering algorithms that are available from most clustering packages.

Here we present hypercluster, a python package and SnakeMake pipeline for parallelized clustering calculations and comparison. The hypercluster package allows users to calculate results from multiple hyperparameters using one or many algorithms, then easily calculate and visualize evaluation metrics for each result (16). The accompanying SnakeMake pipeline allows parallelization on a single computer, across a high performance computing cluster, or on cloud based services (17,18), speeding up optimization, especially for large datasets. In addition, our pipeline has all the advantages of the SnakeMake framework, e.g. easily adding new datasets to analyze, keeping track of progress and simplified bug tracking. Currently, hypercluster can compare all clustering algorithms and evaluation metrics from scikit-learn (19), as well as non-negative matrix factorization (NMF) (20), Louvain and Leiden clustering (21,22). In addition, hypercluster can be extended to employ user-supplied clustering algorithm or evaluation metrics. Given a metric to maximize, hypercluster identifies “best” labels and optionally provides comparisons of labeling results. Even if no single metric can be used to select the best hyperparameters, hypercluster provides several visualizations that help users pick labels by balancing many metrics or picking the most reproducible clusters. Hypercluster provides researchers with a python package and pipeline for flexible, parallelized, distributed and user-friendly algorithm selection and hyper-parameter tuning for unsupervised clustering.

## Design and Implementation

### Requirements and structure

The hypercluster package uses scikit-learn (19), python-igraph (23), leidenalg (24) and louvain-igraph (25) to assign cluster labels and uses scikit-learn and custom metrics to compare clustering algorithms and hyperparameters to find optimal clusters for any given input data (Fig. 1). Hypercluster requires python3, pandas (26), numpy (27), scipy (28), matplotlib (29), seaborn (30), scikit-learn (19), python-igraph (23), leidenalg (24), louvain-igraph (25) and SnakeMake (17). Hypercluster can be run independently of SnakeMake, as a standalone python package. Inputs, outputs and an example workflow are described below, but additional example workflows are provided at https://github.com/ruggleslab/hypercluster/tree/master/examples.

**Fig. 1.**
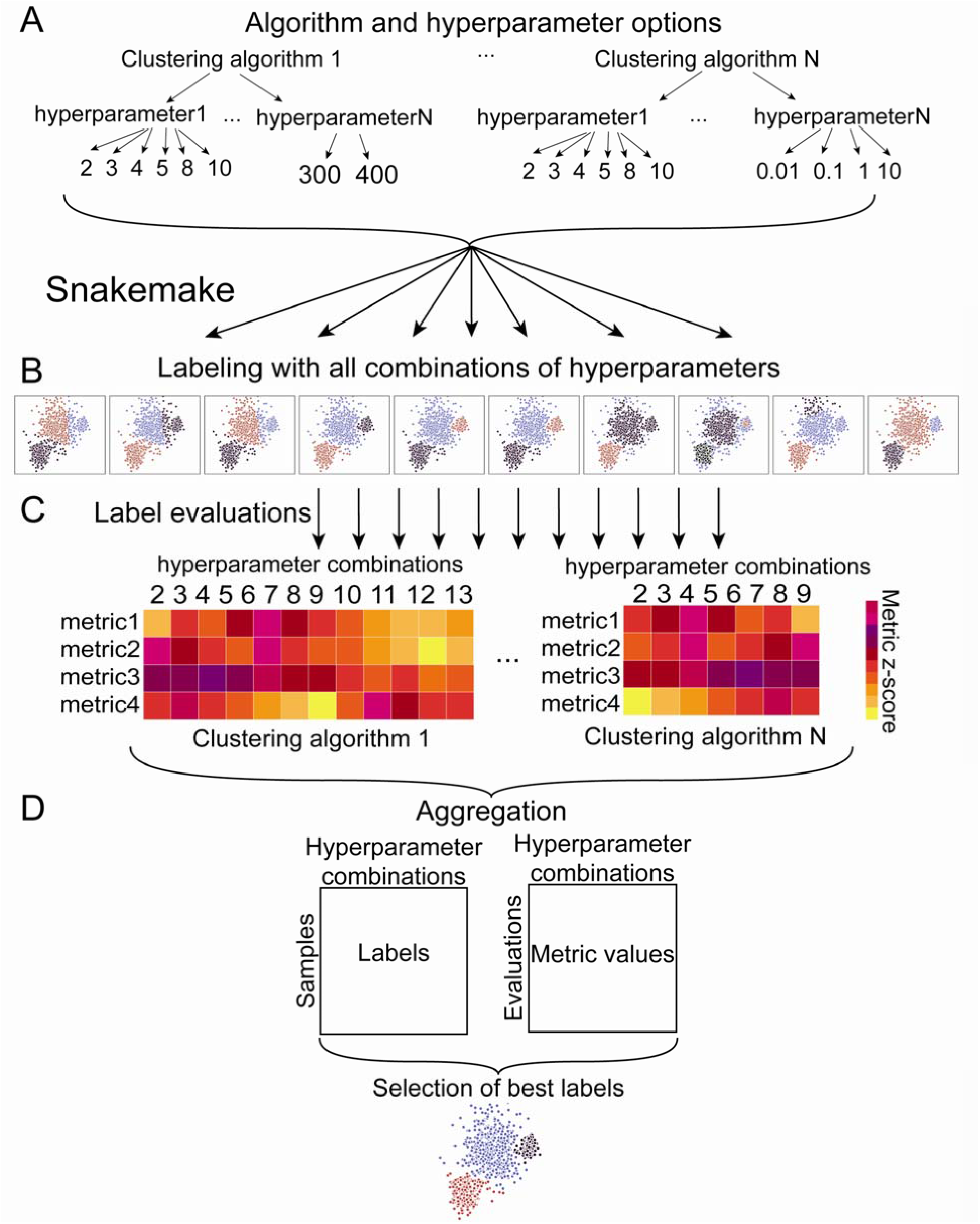
Hypercluster workflow schematic. a) Clustering algorithms and their respective hyperparameters are user-specified. Hypercluster then uses those combinations to create exhaustive configurations, and if selected a random subset is chosen. b) Snakemake is then used to distribute each clustering calculation into different jobs. c) Each set of clustering labels is then evaluated in a separate job by a user-specified list of metrics. d) All clustering results and evaluation results are aggregated into tables. Best labels can also be chosen by a user-specified metric.

### Modes

Hypercluster takes pandas DataFrames as input. For local running, AutoClusterer and MultiAutoClusterer objects can be instantiated with default or user-defined values. To run through hyperparameters for a dataset, users simply provide a pandas DataFrame to the “fit” method on either object. Users evaluate the labeling results by running the “evaluate” method.

### config.yml

SnakeMake allows users to parallelize clustering calculations. To configure the SnakeMake pipeline, users edit a config.yml file (Table 1). In that file, users can specify input and output directories and files (Table 1, lines 1-3, 5-7) and the hyperparameter search space (Fig 1A, Table 1, line 18). Users can specify whether to use exhaustive grid search or random search; if random search is selection, they can specify probability weights for each hyperparameter (Table 1, line 9). Snakemake then schedules performing each clustering algorithm and evaluating the results as a separate job (Fig. 1B). Users can specify which evaluation metrics to apply (Fig. 1C, Table 1, line 10) and add keyword arguments to tune several steps in the process (Table 1, lines 4, 8-9, 11-16). Clustering and evaluation results are then aggregated into final tables (Fig. 1D). Other than the location and names of the input files, everything has a predefined default that allows the pipeline to be used “out of the box.” Users can reference the documentation and examples for more information.

**Table 1.** Parameters in SnakeMake configuration file

| config.yml parameter | Explanation | Example |
| --- | --- | --- |
| 1 input_data_folder | Path to folder in which input data can be found. | /input_data |
| 2 input_data_files | List of prefixes of data files. | ['input_data1', 'input_data2'] |
| 3 gold_standard_file | File name of gold_standard_file, must be in input_data_folder | {'input_data': 'gold_standard_file.txt'} |
| 4 read_csv_kwargs | pandas.read_csv keyword arguments for input data. | {'test_input': {'index_col': [0]}} |
| 5 output_folder | Path to folder into which results should be written. | /results |
| 6 intermediates_folder | Name of subfolder to put intermediate results. | clustering_intermediates |
| 7 clustering_results | Name of subfolder to put aggregated results. | clustering |
| 8 clusterer_kwargs | Additional arguments to pass to clusterers. | KMeans: {'random_state': 8}} |
| 9 generate_parameters_addtl_kwargs | Additional keyword arguments for the hypercluster.AutoClusterer class. | {'KMeans': {'random_search': true}} |
| 10 evaluations | Names of evaluation metrics to use. | ['silhouette_score', 'number_clustered'] |
| 11 eval_kwargs | Additional kwargs per evaluation metric function. | {'silhouette_score': {'random_state': 8}} |
| 12 metric_to_choose_best | Which metric to maximize to choose the labels. | silhouette_score |
| 13 metric_to_compare_labels | Which metric to use to | adjusted_rand_score |
|  | compare label results to each other. |  |
| 14 compare_samples | Whether to made a table and figure with counts of how often each two samples are in the same cluster. | "true" |
| 15 output_kwargs | pandas.to_csv and pandas.read_csv keyword arguments for output tables. | {'evaluations': {'index_col':[0]}, 'labels': {'index_col':[0]}} |
| 16 heatmap_kwargs | Arguments for seaborn.heatmap for pairwise visualizations. | {'vmin':-2, 'vmax':2} |
| 17 optimization_parameters | Which algorithms and corresponding hyperparameters to try. | {'KMeans': {'n_clusters': [5, 6, 7]}} |

**Table 1 Line-by-line explanation of the config.yml for SnakeMake**

### Input data and execution

After specifying the config.yml file, users provide a data table with samples to be clustered as the rows and features as the columns, with the location specified in the config.yml file (Table 1, line2). Users can then simply run “snakemake -s hypercluster.smk --configfile config.yml” in the command line, with any additional SnakeMake flags appropriate for their system. Applying the same configuration to new files or adjusting algorithms and hyperparameter options simply requires editing the config.yml file and rerunning SnakeMake.

### Extending hypercluster

Currently, hypercluster can optimize any clustering algorithm and calculate any evaluation available in scikit-learn (19,31), as well as NMF, Louvain and Leiden clustering. Additional clustering classes and evaluation metric functions can be added by users in the additional_clusterer.py and additional_metrics.py files, respectively, if written to accommodate the same input, outputs and methods (see additional_clusterers.py and additional_metrics.py for examples).

### Outputs

By default, hypercluster outputs a yaml file containing all configurations of the clustering algorithms and hyperparameters that are being searched. For each set of labels, it generates a file containing labels and a file containing evaluations. It also outputs aggregated tables of all labels and evaluations. Finally, given a metric to maximize, hypercluster writes files containing the optimal labels. Optionally, hypercluster will also output a table and heatmap of pairwise comparisons of labeling similarities with a user-specified metric (Figure S1). This figure is particularly useful for finding labels that are robust to differences in hyperparameters. It can also optionally output a table and heatmap showing how often each pair of samples were assigned the same cluster (Figure S2).

## Results

### Unsupervised clustering of RNA-seq on breast cancer patient samples

To illustrate the utility of hypercluster in a disease-relevant context, we applied our method to RNA-seq data from 526 breast cancer patient samples from the Cancer Genome Atlas (TCGA) (32), a dataset that has been previously used for benchmarking clustering algorithms (33). As demonstrated, RNA-seq can be used to classify breast cancer patients into four major PAM50 subtypes (Basal-like, LuminalA, LuminalB, and Her2-enriched), which are based on the expression of 50 specific genes (7,34,35). We removed genes with any missing values and subset to the 500 most variable genes as input for all available algorithms with ranges of hyperparameter conditions. We then compared the sample clustering results from our 500 gene clustering compared with subtypes defined by the PAM50 classifier. This workflow is available on the github examples folder (https://github.com/liliblu/hypercluster/tree/dev/examples).

Hypercluster automatically outputs a visualization of evaluation metrics for all hyperparameter combinations (Fig. 2A), which allows users to quickly see how changing hyperparameters affects clustering result quality. These results highlight how evaluation metrics are not generally convex over ranges of hyperparameters (e.g. silhouette score as n_clusters changes with the KMeans algorithm (Fig. 2A) demonstrating the utility of the exhaustive grid search approach. In addition, our pipeline optionally creates a pairwise comparison of labeling, with a specified user metric (Figure S1) to make it easier to understand how robust and consistent labeling is across algorithms and parameters.

**Fig. 2.**
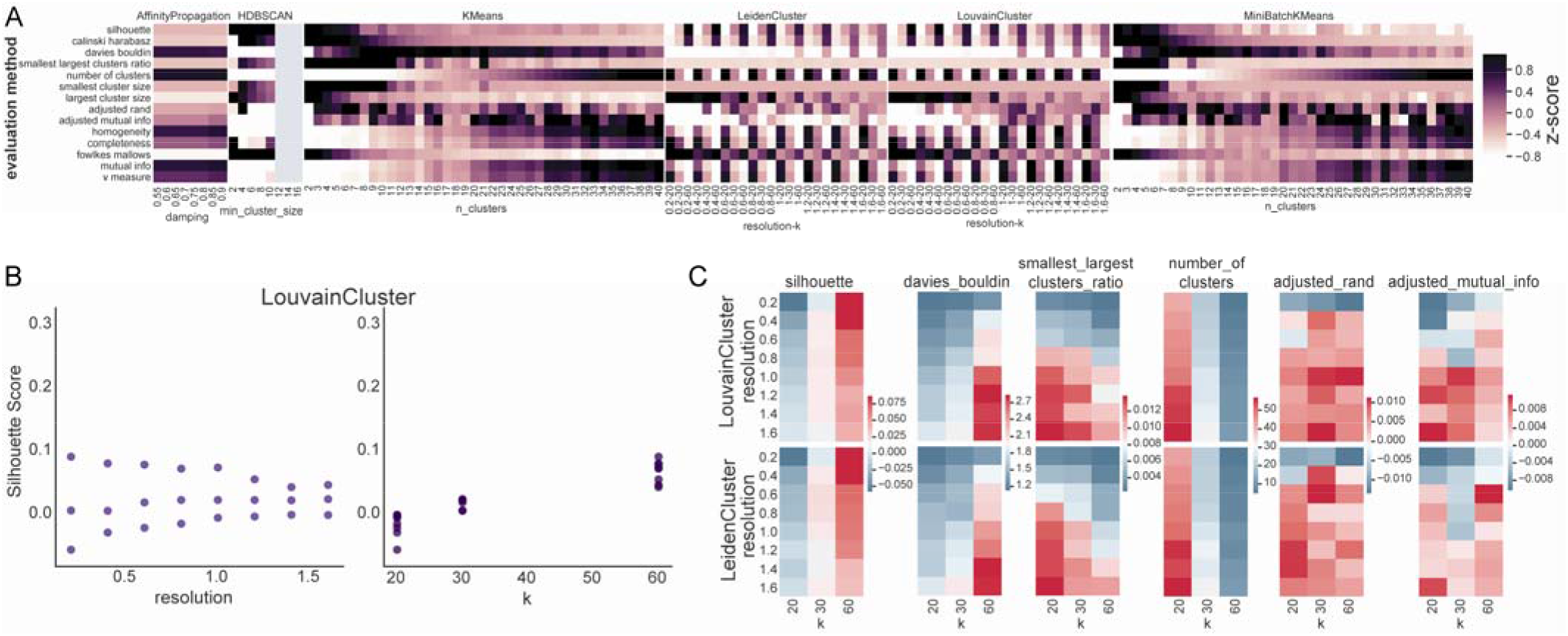
Visualizations of clustering metrics from breast cancer RNA-seq. a) Example automatic output from hypercluster, showing z-scored evaluation values of evaluation metrics for each clustering algorithm and hyperparameter set, including those in 2E. Evaluations applied to clustering from TCGA breast cancer samples. b) Effect of varying resolution (left) and k for shared nearest neighbor matrix (right) on silhouette score, an inherent metric measuring clustering quality, for Louvain clustering. c) Effect of resolution and choice of k on various evaluation metrics for both Louvain (top) and Leiden (bottom) clustering.

Labels and evaluation results are easily accessible for further custom analyses. To demonstrate a possible downstream workflow that hypercluster facilitates, we investigated results from Louvain and Leiden clustering, which are commonly used in scRNA-seq analysis, on the same breast cancer RNA-Seq dataset (36)). Louvain and Leiden clustering are community detection algorithms for networks, usually generated from shared K-nearest-neighbor adjacency matrices. We varied resolution, which affects the number of members in final communities, and the k defining how many nearest neighbors are measured for constructing the adjacency graph (Fig. 2B, C). Resolution and k have significant effects on labeling results and their corresponding evaluations. Interestingly, increasing resolution appears to have opposite effects on clustering quality (e.g. as measured by silhouette score) depending on k, with a large spread of silhouette scores dependent on k at low resolution, converging to similar silhouette scores at higher resolution (Fig. 2B, C). These results highlight the importance of simultaneous tuning of multiple hyperparameters. Plots like those in Fig. 2B, showing the effect of varying each parameter individually on evaluation metrics, can be automatically generated by the visualize_for_picking_best_labels function or listing evaluations in the “screeplot_evals” section of a config.yml file.

To observe if clustering on 500 variable genes can recapitulate PAM50 classification, we identified results that best match PAM50 subtypes according to the adjusted rand score while labeling all samples (Fig. 3). By this metric, the best labels were generated by NMF clustering (37) with n_clusters=4 (Fig. 3A-C). These labels that do diverge from the PAM50 classification correspond to a subset of Luminal A samples that cluster with Luminal B samples (Fig. 3D). Hypercluster allows researchers to compare different algorithms and hyperparameter combinations in a reproducible and convenient way.

**Fig. 3.**
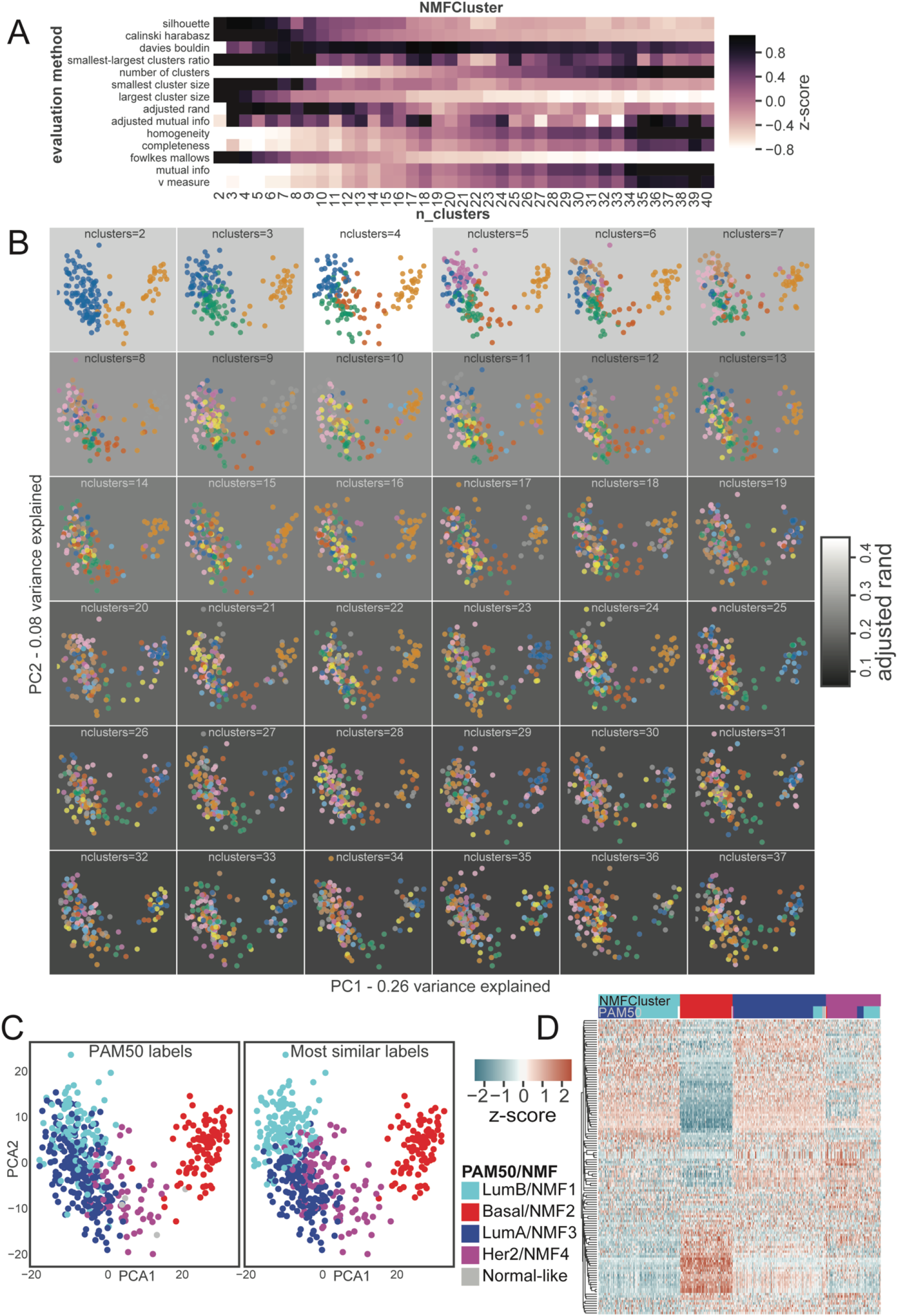
Exploration of NMF clustering results on breast cancer RNA-seq. a) Automatically generated heatmap of evaluation metrics for NMF clustering results. b) PCA projection of 200 random samples colored by labels assigned by NMF clustering. Background color indicates similarity to PAM50 labeling calculated by adjusted rand index. c) PCA projection of samples colored by PAM50 subtypes and most similar NMF clustering labels. d) Heatmap of 125 most variable genes with PAM50 and NMF;n_cluster=4 labels indicated on the top.

### Exploration of bone marrow microenvironment scRNA-seq

To demonstrate hypercluster’s utility for analysis of single cell data sets, we analyzed scRNA-seq from a study investigating the hematopoietic stem cell microenvironment (38) and performed comparative analysis of several clustering algorithms in parallel on a high performance computing cluster utilizing a Slurm scheduler (39). We used normalized expression data from untreated cells sorted for mesenchymal stromal and vascular endothelial, and osteoblast markers, subset to the 2000 most variable genes from the seurat object containing the data (36,38). We then used hypercluster to explore the labeling results from all available clustering algorithms and ranges of relevant hyperparameters. Hypercluster was then used to evaluate labels with every available metric, including metrics that measure inherent labeling quality, as well as comparing new labels to cell types identified in the original study (Fig. 4A). The approach that best recapitulated the published labels was clustering with MiniBatchKMeans with 12 clusters (Fig. 4B-D). These labels differed from published labels largely from swapping cells in the P1 and P2 groups (Fig. 4B), which are both LEPR^+^ subgroups, that were shown to be very similar in the original paper (38). While the original labels were generated using community detection methods like Louvain and Leiden clustering, those methods performed poorly compared to others (Figure S3), likely due to differences in data pre-processing. Varying the number of clusters has variable effects on evaluation results (Fig. 4A, 4C, Figure S3), again highlighting the importance of an exhaustive approach.

**Fig. 4.**
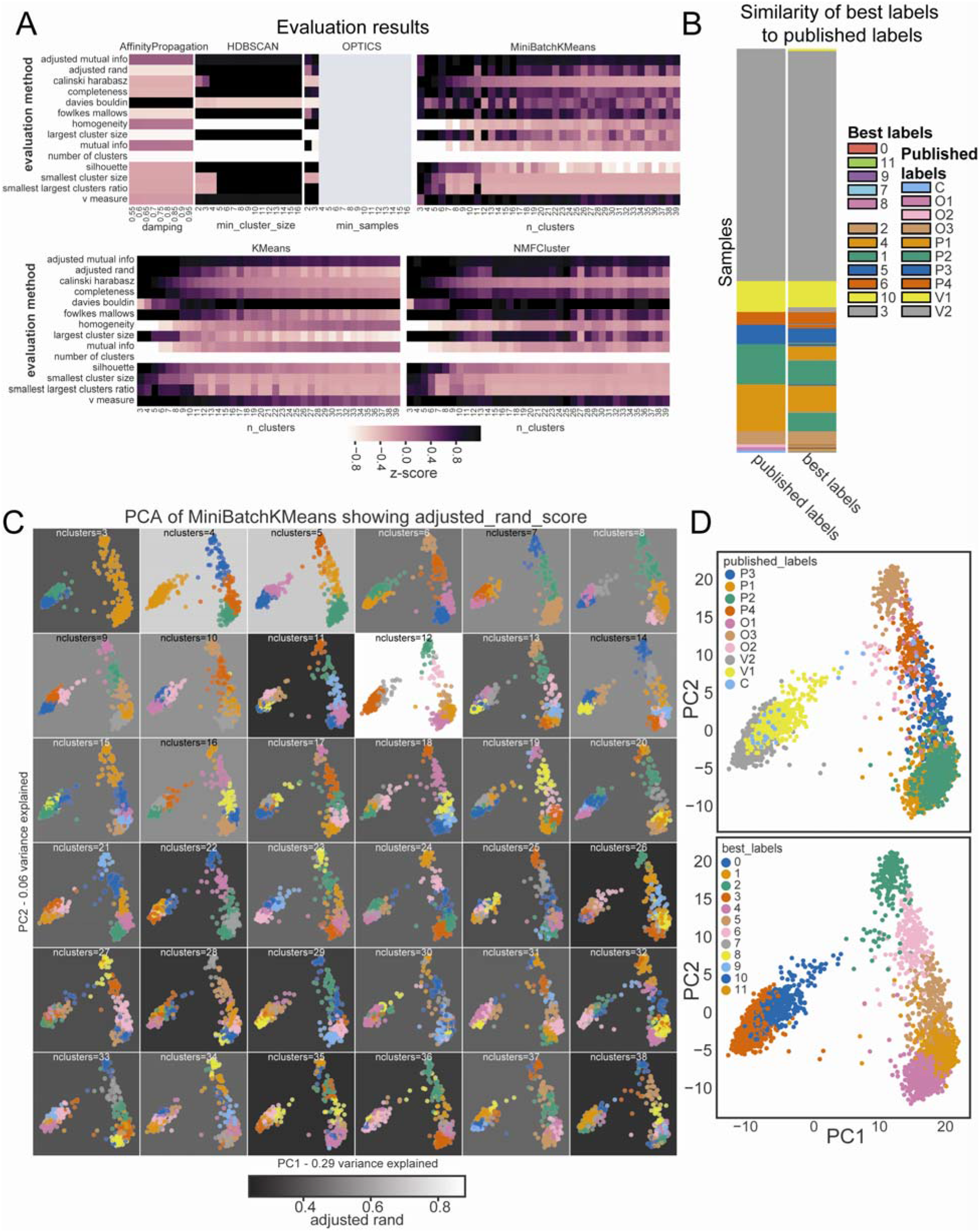
Clustering and evaluation of scRNA-seq data. a) Evaluation metrics for clustering conditions, automatically generated by hypercluster for single cell RNA-seq data. b) Comparison of published labels with best matching calculated labels, MiniBatchKMeans;n_clusters=12. Legend shows mismatched clusters for the best labels on top and clusters with high correspondence to published clusters in the bottom section. c) PCA projection of 700 random cells labeled by MiniBatchKMeans across hyperparameters. d) PCA projection of cells colored by published labels.

## Discussion

Defining groups of molecularly similar patient samples is key to personalizing medical prognosis, diagnosis and treatment strategies, making unsupervised clustering a workhorse for researchers advancing personalized medicine. It is therefore essential that unsupervised clustering is rigorous and not biased by arbitrary hyperparameter selection. While extremely high quality open-source tools such as scikit-learn make unsupervised clustering accessible to many, exhaustively and reproducibly comparing hyperparameters is still challenging; hypercluster solves these issues.

Nearly every step in data analysis pipelines require hyperparameter selection, during which biased or arbitrary parameter selection can greatly impact results. Further, data preprocessing, involving the filtering of datasets to remove low quality or low coverage samples or features (e.g. removing genes with very few reads in RNA-seq), also greatly impacts downstream clustering results. Hypercluster provides a workflow to address the former issue, allowing for comprehensive evaluation of multiple hyperparameters and clustering algorithms simultaneously. The package auto-sklearn (14) provides functionality for automating pre-processing of data tables, which could easily be incorporated upstream of hypercluster to automate the latter. In addition to the simple command line functions, we have also employed SnakeMake for parallelization, a workflow management system already widely used for pipeline optimization (40–46).

If unsupervised clustering is a downstream analytic method of interest, determining which parameters to select can be cumbersome, and possibly inaccurate, without a clustering optimization tool like hypercluster. While it is not always clear how to choose hyperparameters or algorithms in a consistent way (e.g. when two different conditions optimize for different metrics), it is essential to at least understand if the labels one obtains are robust to small changes in algorithm or hyperparameter choice (e.g. as shown in Figure S1). Our package greatly improves the ability of researchers to gain this understanding. In addition to assisting researchers in choosing hyperparameters, hypercluster aids computational biologists who are benchmarking new clustering algorithms, evaluation metrics and pre- or post-processing steps (3). In conclusion, hypercluster streamlines the use of unsupervised clustering to derive biologically relevant structure within data. Most importantly, it eases the prioritization of rigor and reproducibility for researchers using these techniques.

---

**Figure S1.**
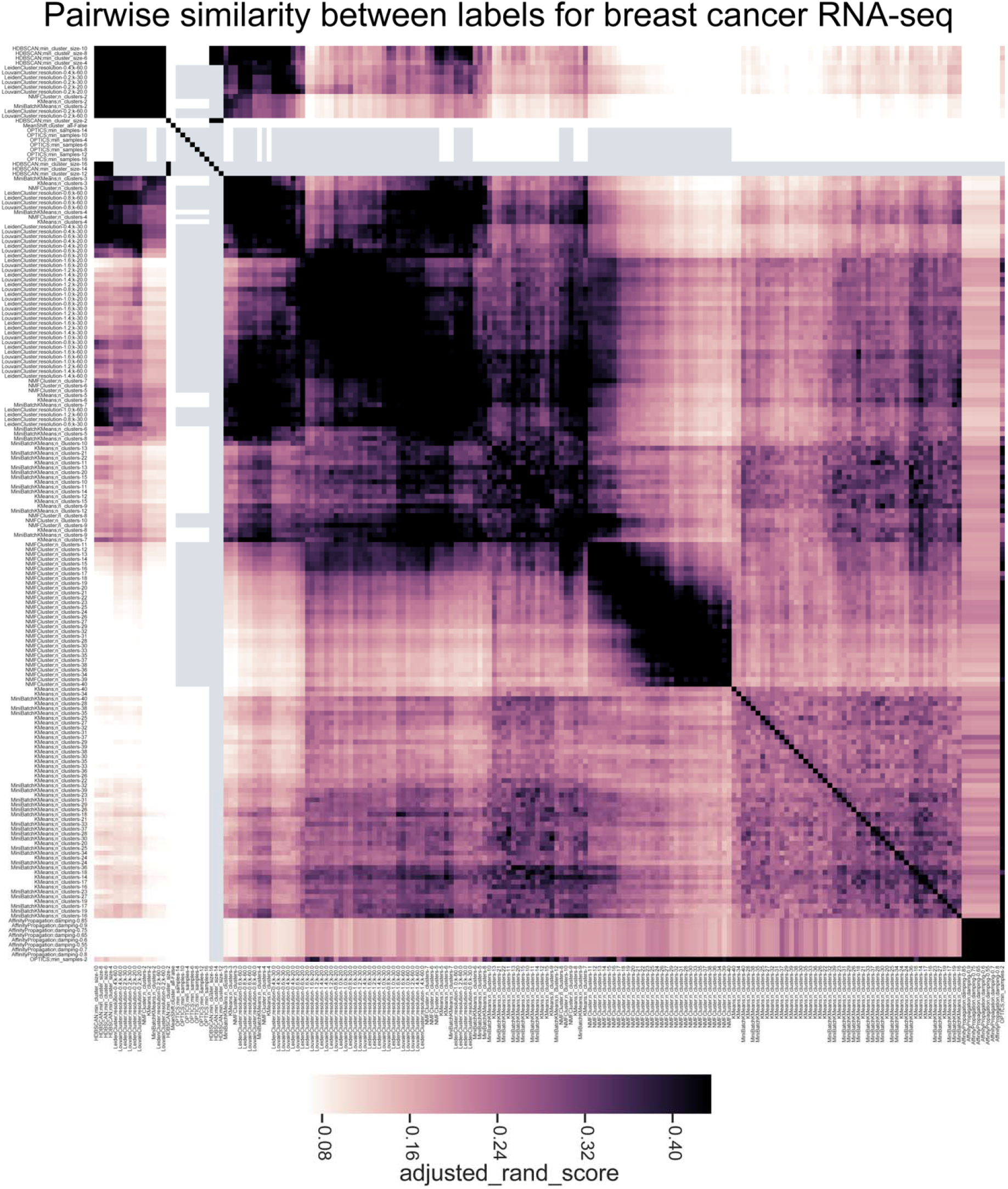
Pairwise label comparisons. Automatically generated heatmap showing pairwise comparison of labeling automatically generated using hypercluster of breast cancer samples. Colors represent adjusted rand index between labels.

**Figure S2.**
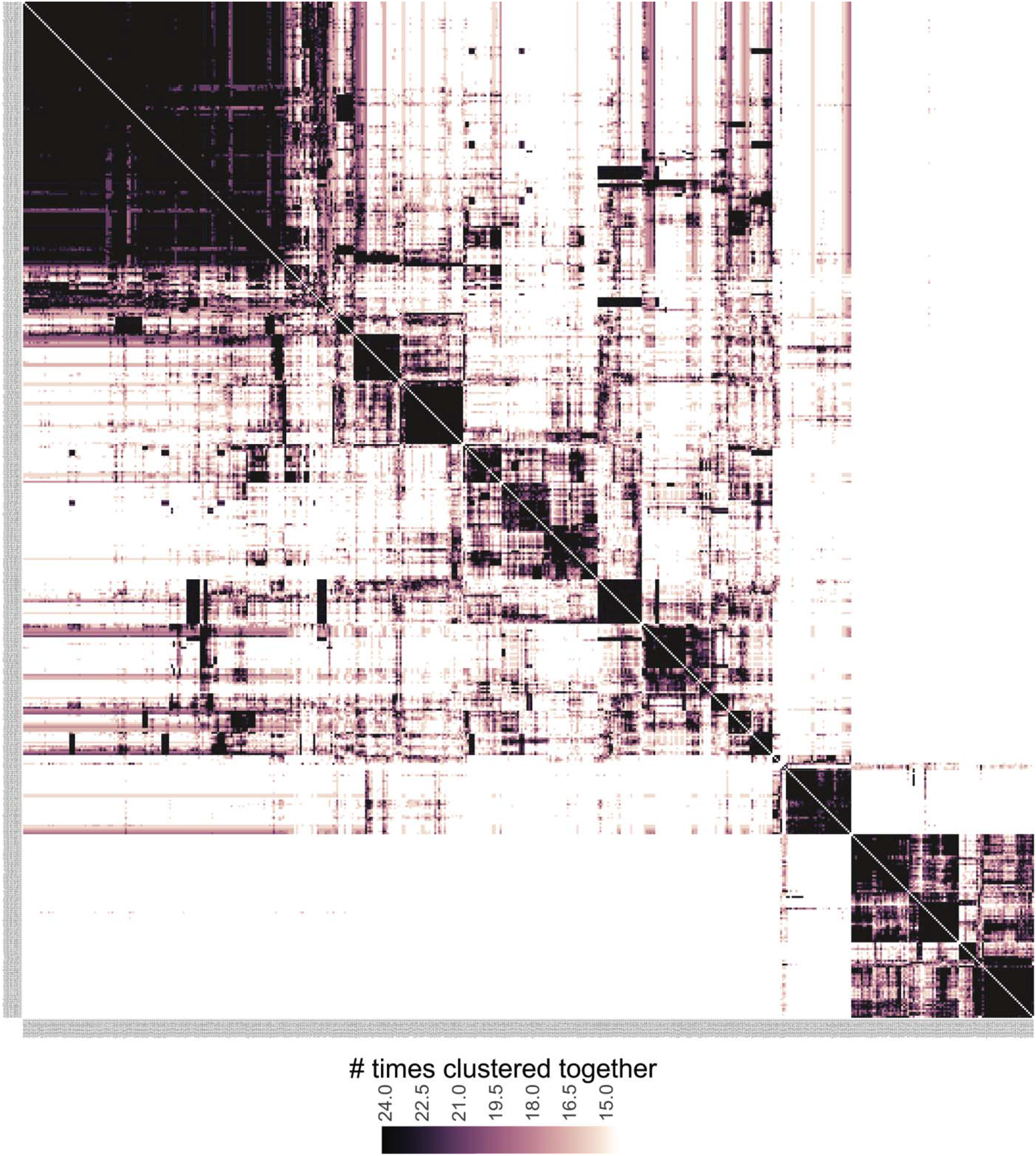
Pairwise sample comparisons. Automatically generated pairwise comparison of breast cancer samples. Color indicates the number of times two samples were assigned the same cluster.

**Figure S3.**
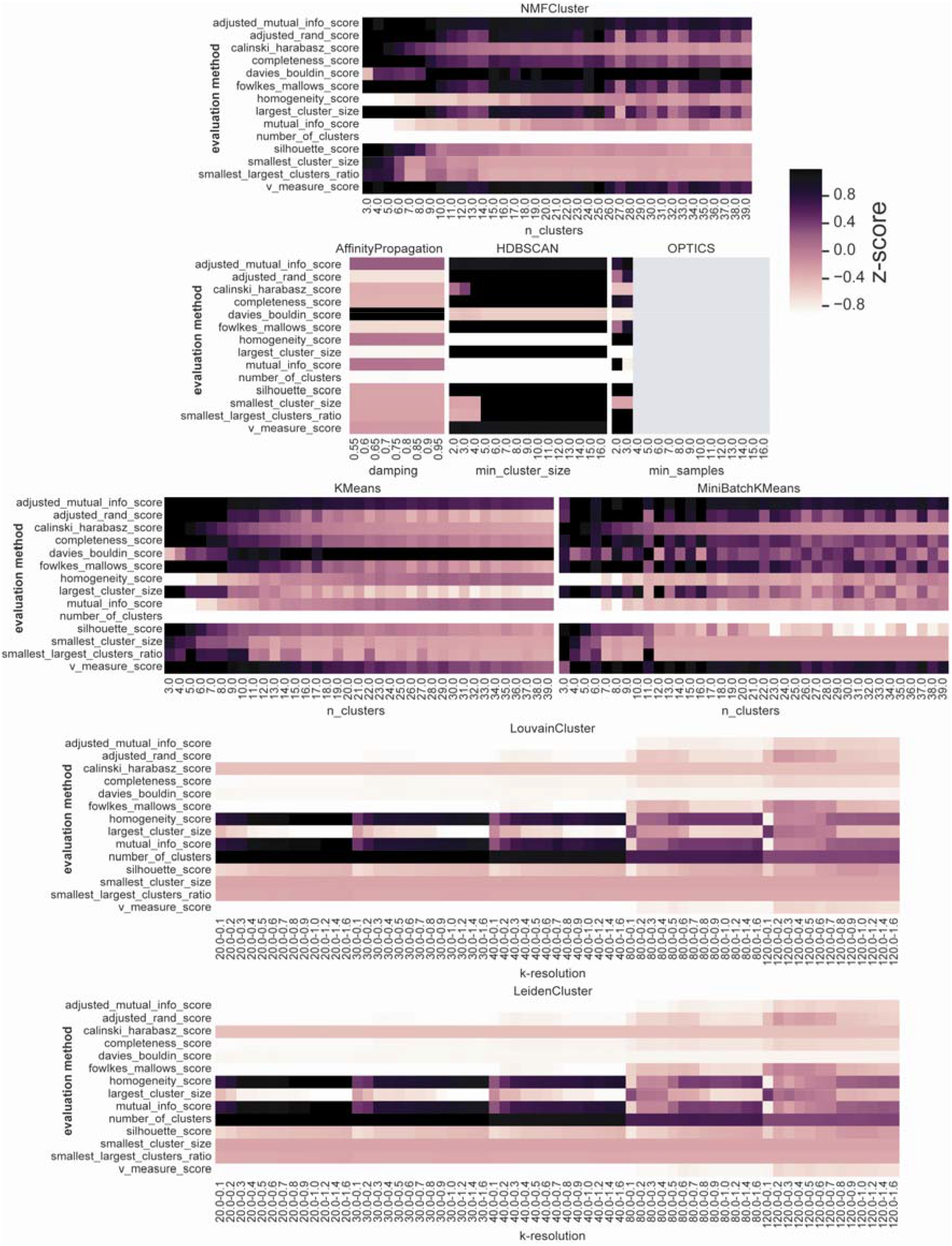
Full evaluations of scRNA-seq clustering. Automatically generated full evaluation metric table from clustering of scRNA-seq stem cell niche cells.

## Acknowledgements

We thank the members of Ruggles and Fenyö labs for their helpful discussions and input. We would like to thank MacIntosh Cornwell for his advice with the SnakeMake pipeline. We would also like to thank Joseph Copper Devlin for his help and advice with implementing Louvain and Leiden clustering.

## Availability of data and materials

Hypercluster is released on pip (pip install hypercluster) and conda (conda install -c bioconda hypercluster). Development versions and installation instructions can be found at our github (https://github.com/liliblu/hypercluster/), tutorials and examples, including all of the code used to create the figures in this paper, can be found here: https://github.com/ruggleslab/hypercluster/tree/master/examples, and documentation can be found here: https://hypercluster.readthedocs.io/en/latest/. Hypercluster is written in python and was developed and tested on MacOS and Linux. Requirements are listed on the github and in the documentation. Hypercluster is available with the MIT licence.

## Authors’ contributions

Conceptualization, Project Administration, Writing: LB and KVR. Data Curation, Formal analysis, Investigation, Methodology, Software, Validation, Visualization: LB. Funding acquisition, Resources, Supervision: KVR.

## Related manuscripts

The authors do not have other related or duplicate manuscripts.

## References

1. Kiselev VY, Andrews TS, Hemberg M. Publisher Correction: Challenges in unsupervised clustering of single-cell RNA-seq data. Nat Rev Genet. 2019 May;20(5):310.

2. Sun S, Zhu J, Ma Y, Zhou X. Accuracy, robustness and scalability of dimensionality reduction methods for single-cell RNA-seq analysis [Internet]. Vol. 20, Genome Biology. 2019. Available from: http://dx.doi.org/10.1186/s13059-019-1898-6

3. Liu X, Song W, Wong BY, Zhang T, Yu S, Lin GN, et al. A comparison framework and guideline of clustering methods for mass cytometry data. Genome Biol. 2019 Dec 23;20(1):297.

4. Parker JS, Mullins M, Cheang MCU, Leung S, Voduc D, Vickery T, et al. Supervised risk predictor of breast cancer based on intrinsic subtypes. J Clin Oncol. 2009 Mar 10;27(8):1160–7.

5. Ohnstad HO, Borgen E, Falk RS, Lien TG, Aaserud M, Sveli MAT, et al. Prognostic value of PAM50 and risk of recurrence score in patients with early-stage breast cancer with long-term follow-up. Breast Cancer Res. 2017 Nov 14;19(1):120.

6. Ali HR, Rueda OM, Chin S-F, Curtis C, Dunning MJ, Aparicio SA, et al. Genome-driven integrated classification of breast cancer validated in over 7,500 samples. Genome Biol. 2014 Aug 28;15(8):431.

7. Perou CM, Sørlie T, Eisen MB, van de Rijn M, Jeffrey SS, Rees CA, et al. Molecular portraits of human breast tumours. Nature. 2000 Aug 17;406(6797):747–52.

8. Capper D, Jones DTW, Sill M, Hovestadt V, Schrimpf D, Sturm D, et al. DNA methylation-based classification of central nervous system tumours. Nature. 2018 Mar 22;555(7697):469–74.

9. Sturm D, Orr BA, Toprak UH, Hovestadt V, Jones DTW, Capper D, et al. New Brain Tumor Entities Emerge from Molecular Classification of CNS-PNETs. Cell. 2016 Feb 25;164(5):1060–72.

10. Hoadley KA, Yau C, Hinoue T, Wolf DM, Lazar AJ, Drill E, et al. Cell-of-Origin Patterns Dominate the Molecular Classification of 10,000 Tumors from 33 Types of Cancer. Cell. 2018 Apr 5;173(2):291–304.e6.

11. Aure MR, Vitelli V, Jernström S, Kumar S, Krohn M, Due EU, et al. Integrative clustering reveals a novel split in the luminal A subtype of breast cancer with impact on outcome. Breast Cancer Res. 2017 Mar 29;19(1):44.

12. Curtis C, Shah SP, Chin S-F, Turashvili G, Rueda OM, Dunning MJ, et al. The genomic and transcriptomic architecture of 2,000 breast tumours reveals novel subgroups. Nature. 2012 Apr 18;486(7403):346–52.

13. Jaskowiak PA, Costa IG, Campello RJGB. Clustering of RNA-Seq samples: Comparison study on cancer data. Methods. 2018 Jan 1;132:42–9.

14. Feurer M, Klein A, Eggensperger K, Springenberg J, Blum M, Hutter F. Efficient and Robust Automated Machine Learning. In: Cortes C, Lawrence ND, Lee DD, Sugiyama M, Garnett R, editors. Advances in Neural Information Processing Systems 28. Curran Associates, Inc.; 2015. p. 2962–70.

15. Barber RF, Ha W. Gradient descent with non-convex constraints: local concavity determines convergence. Inf Inference. 2018 Dec 11;7(4):755–806.

16. Van Craenendonck T, Blockeel H. Using internal validity measures to compare clustering algorithms. Benelearn 2015 Poster presentations (online). 2015;1–8.

17. Köster J, Rahmann S. Snakemake--a scalable bioinformatics workflow engine. Bioinformatics. 2012 Oct 1;28(19):2520–2.

18. Cluster and cloud execution — Snakemake 5.9.1+0.g138720f.dirty documentation [Internet]. [cited 2020 Jan 5]. Available from: https://snakemake.readthedocs.io/en/stable/executing/cluster-cloud.html

19. Pedregosa F, Varoquaux G, Gramfort A, Michel V, Thirion B, Grisel O, et al. Scikit-learn: Machine Learning in Python. J Mach Learn Res. 2011;12(Oct):2825–30.

20. Kim H, Park H. Sparse non-negative matrix factorizations via alternating non-negativity-constrained least squares for microarray data analysis [Internet]. Vol. 23, Bioinformatics. 2007. p. 1495–502. Available from: http://dx.doi.org/10.1093/bioinformatics/btm134

21. Traag VA, Waltman L, van Eck NJ. From Louvain to Leiden: guaranteeing well-connected communities. Sci Rep. 2019 Mar 26;9(1):5233.

22. Traag VA, Krings G, Van Dooren P. Significant scales in community structure. Sci Rep. 2013 Oct 14;3:2930.

23. Csardi G, Nepusz T, Others. The igraph software package for complex network research. InterJournal, complex systems. 2006;1695(5):1–9.

24. Traag V. leidenalg [Internet]. Github; [cited 2020 Jan 27]. Available from: https://github.com/vtraag/leidenalg

25. Traag V. louvain-igraph [Internet]. Github; [cited 2020 Jan 27]. Available from: https://github.com/vtraag/louvain-igraph

26. McKinney W, Others. Data structures for statistical computing in python. In: Proceedings of the 9th Python in Science Conference. Austin, TX; 2010. p. 51–6.

27. Walt S van der, Colbert SC, Varoquaux G. The NumPy Array: A Structure for Efficient Numerical Computation. Comput Sci Eng. 2011 Mar 1;13(2):22–30.

28. Virtanen P, Gommers R, Oliphant TE, Haberland M, Reddy T, Cournapeau D, et al. SciPy 1.0--Fundamental Algorithms for Scientific Computing in Python [Internet]. arXiv [cs.MS]. 2019. Available from: http://arxiv.org/abs/1907.10121

29. Hunter JD. Matplotlib: A 2D Graphics Environment. Comput Sci Eng. 2007 May 1;9(3):90–5.

30. Waskom M, Botvinnik O, O’Kane D, Hobson P, Lukauskas S, Gemperline DC, et al. mwaskom/seaborn: v0.8.1 (September 2017) [Internet]. 2017. Available from: https://zenodo.org/record/883859

31. 2.3. Clustering — scikit-learn 0.22 documentation [Internet]. [cited 2019 Dec 23]. Available from: https://scikit-learn.org/stable/modules/clustering.html

32. Cancer Genome Atlas Network. Comprehensive molecular portraits of human breast tumours. Nature. 2012 Oct 4;490(7418):61–70.

33. Chalise P, Fridley BL. Integrative clustering of multi-level ‘omic data based on non-negative matrix factorization algorithm. PLoS One. 2017 May 1;12(5):e0176278.

34. Sorlie T, Tibshirani R, Parker J, Hastie T, Marron JS, Nobel A, et al. Repeated observation of breast tumor subtypes in independent gene expression data sets. Proc Natl Acad Sci U S A. 2003 Jul 8;100(14):8418–23.

35. Sørlie T, Perou CM, Tibshirani R, Aas T, Geisler S, Johnsen H, et al. Gene expression patterns of breast carcinomas distinguish tumor subclasses with clinical implications. Proc Natl Acad Sci U S A. 2001 Sep 11;98(19):10869–74.

36. Stuart T, Butler A, Hoffman P, Hafemeister C, Papalexi E, Mauck WM 3rd, et al. Comprehensive Integration of Single-Cell Data. Cell. 2019 Jun 13;177(7):1888–902.e21.

37. Tang J, Ceng X, Peng B. New Methods of Data Clustering and Classification Based on NMF [Internet]. 2011 International Conference on Business Computing and Global Informatization. 2011. Available from: http://dx.doi.org/10.1109/bcgin.2011.114

38. Tikhonova AN, Dolgalev I, Hu H, Sivaraj KK, Hoxha E, Cuesta-Domínguez Á, et al. The bone marrow microenvironment at single-cell resolution. Nature. 2019 May;569(7755):222–8.

39. Yoo AB, Jette MA, Grondona M. SLURM: Simple Linux Utility for Resource Management. In: Job Scheduling Strategies for Parallel Processing. Springer Berlin Heidelberg; 2003. p. 44–60.

40. Wang D. hppRNA—a Snakemake-based handy parameter-free pipeline for RNA-Seq analysis of numerous samples. Brief Bioinform. 2018 Jul 20;19(4):622–6.

41. Pranzatelli TJF, Michael DG, Chiorini JA. ATAC2GRN: optimized ATAC-seq and DNase1-seq pipelines for rapid and accurate genome regulatory network inference. BMC Genomics. 2018 Jul 31;19(1):563.

42. Abdelaal T, Michielsen L, Cats D, Hoogduin D, Mei H, Reinders MJT, et al. A comparison of automatic cell identification methods for single-cell RNA sequencing data [Internet]. Vol. 20, Genome Biology. 2019. Available from: http://dx.doi.org/10.1186/s13059-019-1795-z

43. Dirmeier S, Emmenlauer M, Dehio C, Beerenwinkel N. PyBDA: a command line tool for automated analysis of big biological data sets. BMC Bioinformatics. 2019 Nov 12;20(1):564.

44. single-cell-rna-seq [Internet]. Github; [cited 2020 Jan 8]. Available from: https://github.com/snakemake-workflows/single-cell-rna-seq

45. Lun ATL, McCarthy DJ, Marioni JC. A step-by-step workflow for low-level analysis of single-cell RNA-seq data with Bioconductor [Internet]. Vol. 5, F1000Research. 2016. p. 2122. Available from: http://dx.doi.org/10.12688/f1000research.9501.2

46. Soneson C, Robinson MD. Bias, robustness and scalability in single-cell differential expression analysis. Nat Methods. 2018 Apr;15(4):255–61.

